# Multiple particle tracking (MPT) using PEGylated nanoparticles reveals heterogeneity within murine lymph nodes and between lymph nodes at different locations

**DOI:** 10.1101/2022.06.02.494550

**Authors:** Ann Ramirez, Brooke Merwitz, Hannah Lee, Erik Vaughan, Katharina Maisel

**Affiliations:** University of Maryland

**Keywords:** polyethylene glycol (PEG), nanoparticle, microrheology, multiple particle tracking, PEGylation, viscosity, extracellular matrix

## Abstract

Lymph nodes (LNs) are highly structured lymphoid organs that compartmentalize B and T cells in the outer cortex and inner paracortex, respectively, and are supported by a collagen-rich reticular network. Tissue material properties like viscoelasticity and diffusion of materials within extracellular spaces and their implications on cellular behavior and therapeutic delivery have been a recent topic of investigation. Here, we developed a nanoparticle system to investigate the rheological properties, including pore size and viscoelasticity, through multiple particle tracking (MPT) combined with LN slice cultures. Dense coatings with polyethylene glycol (PEG) allow nanoparticles to diffuse within the LN extracellular spaces. Despite differences in function in B and T cell zones, we found that extracellular tissue properties and mesh spacing do not change significantly in the cortex and paracortex, though nanoparticle diffusion was slightly reduced in B cell zones. Interestingly, our data suggest that LN pore sizes are smaller than the previously predicted 10 – 20 μm, with pore sizes ranging from 500 nm - 1.5 μm. Our studies also confirm that LNs exhibit viscoelastic properties, with an initial solid-like response followed by stress-relaxation at higher frequencies. Finally, we found that nanoparticle diffusion is dependent on LN location, with nanoparticles in skin draining LNs exhibiting a higher diffusion coefficient and pore size compared to mesenteric LNs. Our data shed new light onto LN interstitial tissue properties, pore size, and define surface chemistry parameters required for nanoparticles to diffuse within LN interstitium. Our studies also provide both a tool for studying LN interstitium and developing design criteria for nanoparticles targeting LN interstitial spaces.

**Abbreviations:** LNs, FBS, EDC, NHS, ECM, PEG

## 1. Introduction

Lymph nodes (LNs) are found throughout the body and are the key sites where our adaptive immune response is formed. LNs contain an abundance of lymphocytes, or B and T cells, that produce humoral and adaptive immunity, respectively. Antigen presenting cells including macrophages and dendritic cells (DCs) that either reside within the LN cortex and paracortex or migrate from peripheral tissues to the LNs educate lymphocytes by presenting foreign antigens, leading to antigen specific immunity [1]. The structure of the LN is particularly important in maintaining its highly specialized functions. Foreign materials and antigen presenting cells are transported to the LN via afferent lymphatics and into the subcapsular sinus. From here, materials and cells can enter the conduit system that transports them to the LN cortex and paracortex that house B and T cells, respectively. The intricate extracellular structure of the LN, including the conduit system, which consists of vessel-like structures, and supporting extracellular matrix (ECM) have been a topic of study in the last two decades.

Structure-function relationships, including mechanical properties of the LN as a result of its extracellular material and conduit composition throughout inflammatory processes, and their effect on development of the immune response, have been more recently elucidated [2–6]. During homeostasis, LN interstitial spaces are thought to range between 10 to 20 μm, allowing cells to freely move while adhering to ECM fibers and conduits [7]. However, in times of rapid LN growth (i.e., inflammation), these spaces increase in size while ECM and conduit density is reduced [2]. Some research suggests that flow within the conduit network is interrupted during expansion and that the density of matrix components, including collagen I, IV, and VI, decreases during inflammation [2]. Additionally, matrix components begin to fragment during expansion, leaving cellular components to bear the increased tension in the expanded tissue [8]. This tissue remodelling and accumulation of macromolecules during inflammatory processes are likely to affect the interstitial viscosity and viscoelasticity of the interstitial tissue space [2, 6, 9]. Recent evidence has demonstrated that tissue viscoelasticity and fluid viscosity can affect a variety of cellular behaviors, including adhesion to extracellular matrix and migration [10–12]. However, the difficulty in accessing LNs due to their small size in animal models has made it difficult to study LN interstitial viscosity and viscoelasticity until now.

Here, we have combined two methods that have emerged from engineering and biology to study interstitial tissue spaces: ex vivo tissue slice cultures and microrheology using multiple particle tracking (MPT). Ex vivo tissue slice cultures were first developed to study brain tissue pathogenesis and have since evolved to be used for development of brain and cancer therapeutics, and more recently to study cell migration and material diffusion throughout the LNs [13, 14]. Tissue slices are particularly physiologically relevant; the tissue architecture stays intact, thus allowing for spatial and temporal resolution of cellular behaviors within their physiologically relevant microenvironment[15].

MPT is a technique used to study the Brownian motion of particles within different materials. This method was originally developed to study properties of polymeric materials, and more recently was applied to studying biological hydrogels, such as mucus and ECM [16]. Video capturing of nanoparticle diffusion and subsequent analysis allows the calculation of mean squared displacement (MSD) of the nanoparticles and from this, various properties can be extrapolated [16]. This method has also been coined ‘microrheology,’ as it allows calculation of material properties like diffusion coefficient, viscosity, pore size, elasticity, and plasticity that normally require larger volumes of materials to be placed into a rheometer (which is not always feasible, as is the case with LN tissues).

Using MPT to characterize interstitial tissues requires a nanoparticle probe system that minimally interacts with, or binds to, the biological environment. Polyethylene glycol (PEG) has previously been shown to reduce nanoparticle interaction with biological materials, particularly when PEG coatings on nanoparticles are dense [17–19]. In this study, we used 100 and 200 nm polystyrene (PS) core nanoparticles to determine how PEG density affects nanoparticle interaction with LN extracellular environments. We then used MPT with the newly designed nanoparticle probe to assess microrheological properties of LN interstitial tissue, both in the different regions within LNs and in LNs from different regions in the body.

## 2. Materials and methods

### Tissue collection

Female C57BL/6 mice 8-20 weeks old (Jackson Laboratory) were euthanized by CO2 inhalation. Inguinal, brachial, and mesenteric LNs were collected, fat surrounding the LNs was removed, and LNs were placed in cold 1X PBS supplemented with 2% fetal bovine serum (FBS, Sigma). All animal work was approved by the Institutional Animal Care and Use Committee at the University of Maryland, College Park.

### LN slicing

LNs were embedded in a 6% w/v low-melting agarose (Thomas Scientific) using 35 mm Petri dishes. LNs were placed into the agarose with the wide side of the node facing up and a 10 mm biopsy punch (Robbins Instruments) was used to punch out the embedded tissues. The gel pieces were oriented so that the tissue was at the top of the gel piece and glued (CA 221 fast curing glue, Best Klebstoffe) onto the specimen holder of a Leica VT1000S vibratome. The Leica VT1000S vibratome was set to a speed of 3.9mm/s, frequency set to 0.3Hz, and amplitude set to 0.6mm and up to 4 LNs were sliced simultaneously. 1X PBS was placed into the buffer tray and ice was continually placed in the area surrounding the buffer tray to ensure the 1X PBS was cold throughout the duration of the slicing. LN slices were collected using a brush and placed into RPMI (Thermo Fisher) supplemented with 10% FBS (Sigma), 1× L-glutamine (Corning), 50 U/mL Pen/Strep (Sigma), 50 μM beta-mercaptoethanol (Thermo Fisher)), 1 mM sodium pyruvate (Thermo Fisher), 1× nonessential amino acids (Fisher Scientific), and 20 mM HEPES (GE Healthcare). LN slices were incubated at 37 °C with 5% CO_2_ in media for at least 1h prior to performing multiple particle tracking.

### LN staining

Slices were placed on a microscopy slide and Fc receptors on cells were blocked to avoid nonspecific staining using 20 μl purified anti-mouse CD16/32 antibody (BioLegend) in 1X PBS with 2% FBS at a concentration of 20μg/ml for 30 min at 37 °C. To stain for B cells, slices were incubated with 20 μg/ml anti-B220 antibodies (AF647-αB220 Rat IgG2a, κ or FITC-αB220 Rat IgG2a, κ, BioLegend) at 37 °C for 1 h. To stain for T cells, slices were incubated with 20 μg/ml anti-CD45 (Pac-Blue-αCD45 Rat IgG2a, κ, BioLegend). Slices were washed 3x with 1X PBS for 10 min per wash.

### Nanoparticle formulation

To formulate nanoparticles for MPT, we PEGylated 100, 200, or 500 nm carboxylate-modified polystyrene microspheres (Fluospheres™, Thermo Fisher) with 5 kDa PEG using EDC/NHS chemistry [20]. Briefly, particles were dissolved at 0.1 % w/v in 200 mM borate buffer. Amine terminated 5 kDa PEG (Creative PEGworks) was conjugated to the particle surface using 0.02 mM ethyl-3-(3-dimethylaminopropyl) carbodiimide (EDC) and 7 mM N-hydroxysuccinimide (NHS) and left to react for at least 4 hours at room temperature in a rotator and protected from light. Theoretical 100% PEG coverage to carboxylate groups was achieved with the following PEG concentrations: 350 μM, 90 μM, and 310 μM for 100, 200, and 500 nm particles, respectively. Theoretical 5% PEG coverage (mushroom conformation) was achieved with the following PEG concentrations: 17.4 μM, 4.5 μM, and 15.5 μM for 100, 200, and 500 nm particles, respectively. Particles were washed 3 times in deionized (DI) water using a 100 kDa filter (Amicon Ultra-0.5 Centrifugal Filter Unit) in a centrifuge at 17g for 12 minutes. Particles were resuspended in DI water at the same concentration as the package formulation. Particle size and ζ-potential was characterized using dynamic light scattering (DLS).

### Nanoparticle tracking

Up to 2 μl of a 1:1000 nanoparticle solution was placed on top of each LN slice. A coverslip was placed on the slice and stayed on by means of capillary forces. Movies were captured using a 63X objective on a ZEISS Axioscope 5 microscope and by using Zeiss software (ZEN lite) at a temporal resolution of 30 or 50 ms for 20 s. Mean squared displacement (MSD) and nanoparticle trajectory was calculated using MATLAB, with a minimum of 10 frames for each particle [21]. MSD was calculated using 〈Δ*r*^2^(τ)〉 = [*x(t + τ)* − *x(t)*]^2^ + [*y*(*t* + τ) − *y(t)*]^2^ with at least 70 particles.

### Statistics

Group analysis was performed using a 2-way ANOVA. Unpaired Student’s t-test was used to examine differences between only two groups. A value of *p* < 0.05 was considered significant (GraphPad). All data is presented as mean ± standard error of the mean (SEM).

## 3. Results and discussion

### Dense PEG coatings enhance nanoparticle diffusion through LN extracellular tissue spaces

Nanoparticles have emerged as biophysical tools to study rheological properties of biological materials, particularly through the development of MPT. Size of particles is an important consideration to take when MPT is being used to extrapolate values like pore size. Typically, particle size is chosen to be much larger than the mesh size in the sample [16]. This ensures that the bulk area is being measured and not just the particle environment. We chose to pick a particle size smaller than the predicted mesh size to emphasize the heterogeneity of the LN ECM space. Additionally, using MPT for determining material properties like microrheology of a biological sample requires the nanoparticles to minimally interact with the native environment, as charge or hydrophilicity-based interactions can affect microrheological properties calculated via MPT [16]. PEGylation has been used to design nanoparticles that are able to cross biological barriers, including cellular barriers [22–24] and extracellular tissue structures such as mucus [21, 25–27] and brain/tumor ECM [28–30]. A key parameter to minimize nanoparticle-biological material interactions is the density of PEG on the nanoparticle surface [17]. To test how PEG density affects nanoparticle interactions with LN interstitial tissue, we formulated 100 nm or 200 nm core PS nanoparticles with either a low density of PEG, often referred to as ‘mushroom’ conformation in literature, or a high density of PEG, referred to as ‘dense brush’ (**Table 1**). The ratio between the PEG chain’s Flory radius (Rf, the theoretical space taken up by an unconstrained linear molecule) and the grafting distance (D, distance between each PEG chain) on the surface of the NP determines the conformation of the PEG chain [17]. The Flory radius is determined by *RF* ~ *a*N^3/5^, where *a* is the monomer length and N is the degree of polymerization [17]. Therefore, PEG chain arrangements with Rf/D values < 1 are considered to be in a mushroom conformation, while those with Rf/D values > 3-4 are considered to be in a dense brush conformation. Non-coated PS nanoparticles had a highly negative surface ζ-potential, due to functionalization with carboxylic acid groups (**Table 1**). Increasing amounts of PEG on nanoparticles reduced the surface ζ-potential to near neutral charge (**Table 1**) and we confirmed that nanoparticles were coated either with mushroom (Rf/D close to 1) or dense (Rf/D > 4) PEG conformations (**Table 1**) [23].

**Table 1.** Characterization of different densities of PEG on polystyrene beads. Size, zeta potential, and Rf/D of 100, 200, and 500 nm nanoparticles. Size and zeta potential were determined using dynamic light scattering (DLS). Data shown as mean ± SEM (*n* = 1 - 6).

| | Size (nm) | ζ-potential (mV) | R<sub>f</sub>/D |
| --- | --- | --- | --- |
| 100 nm PS | 110 ± 0.6 | - 17.4 ± 3.5 | - |
| 200 nm PS | 200 ± 1.0 | - 40.0 ± 2.0 | - |
| 100 nm PSPEG<sub>M</sub> | 130 ± 1.6 | - 6.0 ± 2.9 | 0.2 ± 0.0 |
| 200 nm PSPEG<sub>M</sub> | 211 ± 1.1 | - 9.9 ± 0.5 | 1.5 ± 0.1 |
| 100 nm PSPEG<sub>D</sub> | 130 ± 0.8 | -0.3 ± 0.1 | 3.4 ± 0.3 |
| 200 nm PSPEG<sub>D</sub> | 220 ± 2.4 | - 5.0 ± 0.5 | 5.0 ± 0.8 |
| 500 nm PSPEG<sub>D</sub> | 520 ± 2.0 | -6.4 ± 0.4 | 3.6 ± 0.0 |

Next, we assessed the diffusion of PS and PSPEG nanoparticles in LN slices ex vivo. We found that uncoated PS nanoparticles (PS) minimally diffused through LN interstitial tissue, as indicated by their low MSD and diffusion coefficient, D (**Figure 1A, B**) with D = 0.2 ± 0.1 μm^2^/s and 0.3 ± 0.2 μm^2^/s for 100 nm and 200 nm non-PEG coated nanoparticles, respectively. Low density PEG (PSPEG_M_) slightly increased nanoparticle MSD of both 100 and 200 nm nanoparticles, but diffusion remained low (**Figure 1A, B**) compared to densely coated nanoparticles (PSPEG_D_), with diffusion coefficients of 0.7 ± 0.4 μm^2^/s and 0.8 ± 0.4 μm^2^/s for 100 nm and 200 nm nanoparticles, respectively. Dense PEG coatings led to an average mean diffusion coefficient of 3 ± 0.8 μm^2^/s for 100 nm and 2.8 ± 0.8 μm^2^/s for 200 nm nanoparticles, which are close to the theoretical diffusion coefficient of these nanoparticles in water, ~4.3 μm^2^/s for 100 nm and ~2.15 μm^2^/s for 200 nm nanoparticles (**Figure 1B**). This suggests that densely PEGylated nanoparticles are diffusing through LN interstitial tissue without significant interactions with ECM materials or cells. Representative trajectories of each particle type and size also suggest that densely PEGylated nanoparticles diffused the most effectively (**Figure 1C**). Distribution of nanoparticles’ MSDs demonstrates that more of both the PSPEG_M_ and PS have MSDs smaller than their size, compared to PSPEG_D_ (**Figure 1D**).

**Figure 1:**
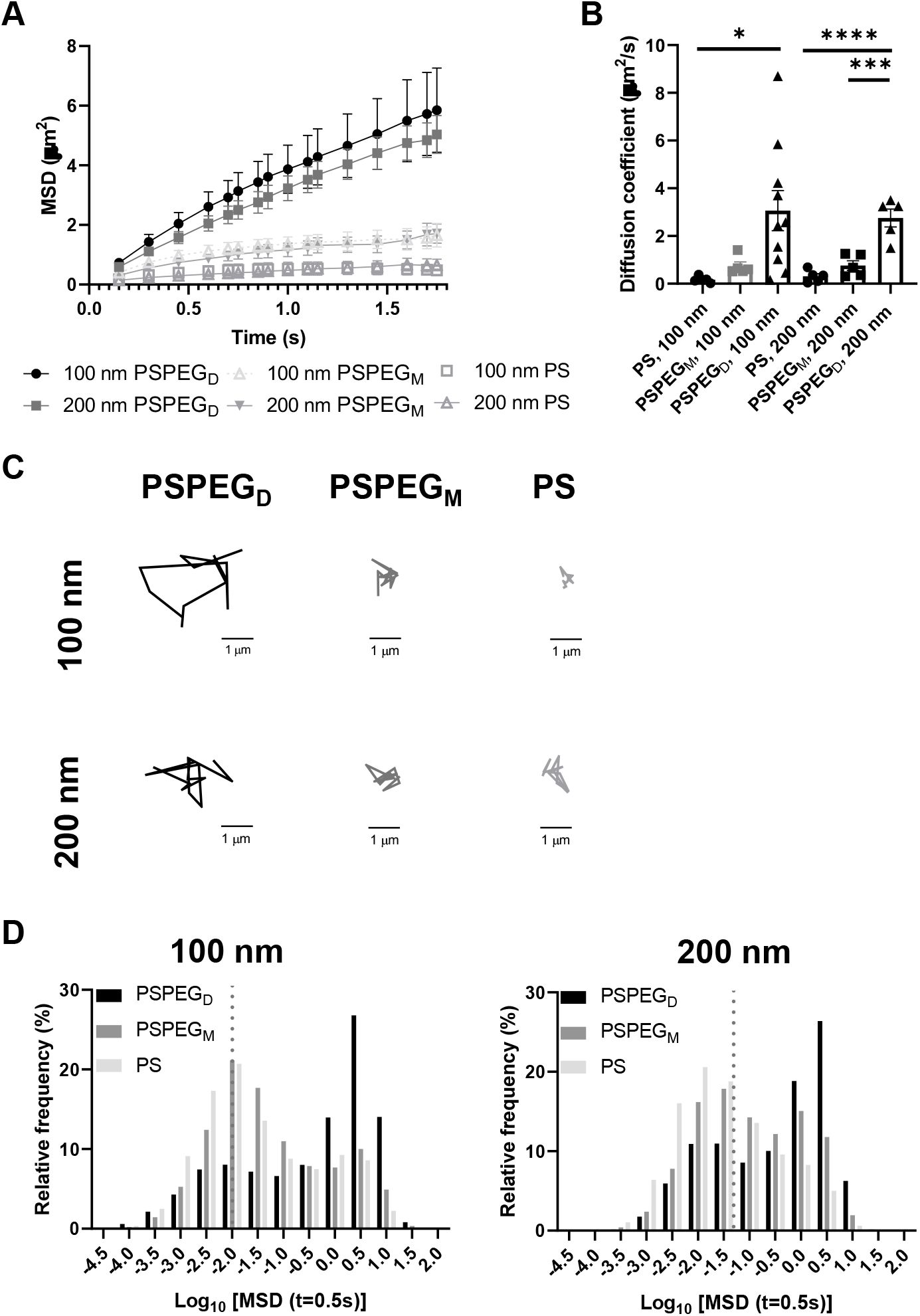
Dense PEGylation is required for particle diffusion in sdLN and mLN tissue slices. **(A)** Mean squared displacement (MSD) and **(B)** diffusion coefficient ± SEM of non-PEGylated, low-density PEGylated, and high-density PEGylated 100nm and 200nm nanoparticles **(C)** Trajectories of representative particles. **(D)** Frequency distribution of 100 and 200 nm particles at different densities. Diffusivity values to the left of the dotted line indicate particles with MSD values less than the particle diameter. Data shown as mean ± SEM (*n* = 5 - 10).

Prior literature has shown that nanoparticle material properties like shape, size, and charge can modulate their interactions with ECM barriers. To be able to cross extracellular barriers and reach cells within a tissue, a small size and non-‘sticky’ surface chemistry are required [31]. Several studies have suggested that nanoparticles with charge similar to that of the ECM materials can enhance nanoparticle diffusion through these barriers due to repulsive forces [32–36]. Having opposing charges, in contrast, usually leads nanoparticles to become stuck in the extracellular matrix spaces [32–36]. Neutral nanoparticle charge, like those used in our studies, has also been suggested to improve nanoparticle penetration through ECM [29, 34, 36]. Additionally, some studies have determined the effect of PEG density on nanoparticle diffusion in ECM. For instance, Nance et al. showed that dense PEG coatings are required for nanoparticle diffusion through brain parenchyma [29]. Additionally, researchers have found that increasing PEG density enhances liposome and polymeric nanoparticle diffusion in collagen gels [30, 37]. Our data recapitulate these findings and suggest that dense PEGylation is required for nanoparticles to effectively diffuse through LN ECM spaces.

### MPT reveals similar pore size and viscosity within different regions of the LN

LN structure and function are closely related. B cells form follicles in the outer cortex of the LN, while T cells are mostly found in the inner paracortex (**Figure 2A–B**). The reticular network spans throughout the LN and is more densely packed in the T cell zone (**Figure 2A–B**). Podoplanin (PDPN) which is used to define the conduit network, is more densely found within the T cell one compared to the B cell zone (**Figure 2C**) [2]. Typically, only materials less than 70 kDa can enter the reticular network to cross into the cortex and paracortex [38]. Recent work suggests that lymphatic vessels are also more concentrated in the T cell zone, further supporting the hypothesis that the T cell zone is more densely packed than the B cell zone [39]. We next investigated interstitial tissue in the T and B cell zones using our newly developed nanoparticle probes. We first confirmed that PSPEG_D_ diffused effectively within B and T cell zones, while PSPEG_M_ and PS did not (**Supplementary Figure 2**). These findings are consistent both with our earlier findings and with previous literature that has suggested that dense PEGylation is required to maximize nanoparticle diffusion through ECM [37]. When we specifically compare the diffusive nanoparticle MSD between T and B cell zones, we found that there is no significant difference based on region within the LN for 100 nm, 200 nm, and 500 nm PSPEG_D_ (**Figure 2D**). However, there are significant differences in the B cell zone between 100 nm and 200 nm PSPEG_D_ particles at times below 1.3 ms. Similarly, there are significant differences between 200 nm and 500 nm at times below 1.7 ms. MSD in the T cell zone were only significantly different between 200 nm and 500 nm PSPEG_D_ particles at times between 1.15 ms and 1.6 ms. We observed that 500 nm PSPEG_D_ had significantly lower MSD in both the B and T cell zones, indicating these spaces in LNs restrict movement of larger particles. However, we did find that nanoparticles had slightly lower diffusion in T compared to B cell zones for all PSPEG_D_ (**Figure 2D**), which is also reflected in the distribution of nanoparticle MSDs within the B vs. T cell zones (**Figure 2E**). We found that both 100 nm and 200 nm PSPEG_D_ have diffusion coefficients similar to that in water (see earlier, **Figure 2G**), suggesting that PSPEG_D_ are freely diffusing in both B and T cell zones.

**Figure 2:**
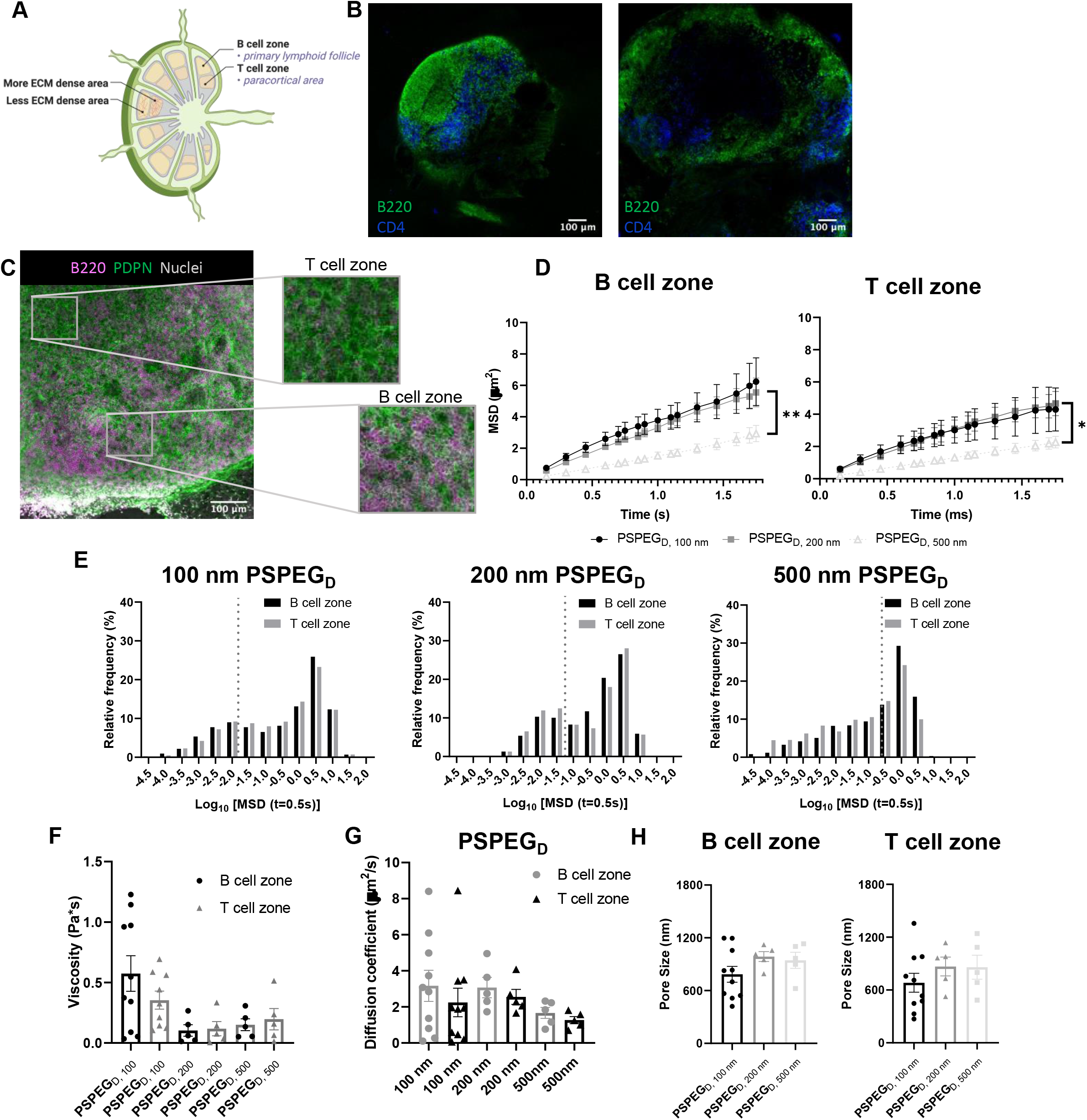
Diffusion does not change in either the B or T cell zone. **(A)** Cross-sectional view of the different zones within a LN slice. **(B)** Confocal images of the first 0-300 μm (left) and second 300-600 μm (right) (B cells – cyan, T cells – blue) slice of a sdLN. **(C)** Cross-sectional view of mLN with stained Podoplanin (PDPN, green) and B cells (magenta). Zoomed in view shows change in PDPN density between T and B cell zones. **(D)** MSD of high-density PEGylated 100, 200, and 500 nm nanoparticles in B or T cell zones of sdLN and mLNs. **(E)** Frequency distribution (diffusivity values to the left of the dotted line indicate particles with MSD values less than the particle diameter), **(F)** viscosity, **(G)** diffusion coefficient, and **(H)** pore size of densely PEGylated 100, 200, and 500 nm particles in B and T cell zones of sdLN and mLNs. Data shown as mean ± SEM (*n* = 5 - 10).

Although not significant, the diffusion coefficient of particles in the T cell zone is 1.5-fold lower than that in the B cell zone, with 2.2 ± 0.8 μm^2^/s compared to 3.2 ± 0.9 μm^2^/s, respectively for 100 nm PSPEG_D_ (**Figure 2G**). The densely packed nature of the T cell zone, particularly due to reticular fibers, has been hypothesized to stiffen this region [38] and may be the reason for the decrease in the diffusion coefficient. In addition to reticular fiber density, the viscosity of the fluid within the ECM may also have a role in the differences in diffusivity. We have found that viscosity in the T and B cell zones is similar, though viscosity in the B cell zones is slightly higher at ± 0.1 Pa∙s compared to 0.4 ± 0.07 Pa∙s in the T cell zone for 100 nm PSPEG_D_ (**Figure 2F**). Studies have suggested that cell speed increases in fluids with higher viscosity [40, 41]. Recent studies have found that T cell motility changes throughout the LN, and is potentially dependent on the region within which the T cell is found [42]. Interestingly, we have also observed that diffusion of nanoparticles may change throughout the depth of the LNs (**Supplementary Figure 1**), suggesting that viscosity may also change throughout the LN. It is possible that extracellular fluid viscosity has an additional role in modulating cell motility and migration throughout the LN. However, viscosity of the LN interstitial fluid is still not well studied, and many factors can influence viscosity including extracellular remodeling or secretion of proteins and other macromolecules [2, 6, 9], and all these factors may be further affected by states such as inflammation. More research is needed to fully understand how material properties like viscosity change within the LN environment and how this may affect cell behavior within the LNs.

From the MPT data, we can extract information about nanoparticle diffusion and use this to extrapolate mesh spacing or pore size of the material (see Materials and Methods). We found that the extracellular spacing based on MPT is close to 0.5 – 1 μm (**Figure 2H**), and pore size is highly variable, as indicated by the range of values observed (**Figure 2H**). Our findings are in contrast to prior literature that suggests that the mesh spacing or pore size of the LN is closer to 10-20 μm, at least one order of magnitude above our estimated pore size [43]. However, mesh spacing literature focuses only on the pore size or mesh spacing based on ECM material and does not account for space taken up by cells, themselves. We expect that the difference in mesh spacing observed through MPT is likely due to the high density of cells found within the LNs. During inflammatory processes, such as disease, infection, or vaccination, the LN significantly expands and can grow to be more than 10-fold its original size and weight. We expect that inflammation would further affect pore size, due to changes in the ECM and reticular fibers, and the recruitment of additional immune cells [44]. How these factors affect extracellular tissue mesh spacing or pore size, as well as nanoparticle diffusivity and viscosity, is currently under investigation in our lab.

### Microrheology confirms that LNs are viscoelastic

MPT has emerged as a tool to determine pore size and viscosity of hydrogels, and as a rheological tool to assess viscoelastic properties of biomaterials. We used our nanoparticle tracking data and the Stokes-Einstein equations to determine viscoelasticity of LN interstitial tissue. Several tissues have previously been shown to behave as viscoelastic materials, exhibiting a solid-like response, but over time there is stress relaxation [45]. This behavior can be characterized using elastic (G’) and loss (G’’) moduli, where G’ values describe elastic properties of a material, while G” values describe viscous properties. We found that in 100 nm, 200 nm, and 500 nm PSPEG_D_ in either zone, the tissue experiences a slight decrease in G’ and G” at lower frequencies (< 0.7 Hz) and a slight increase in G’ and G” at frequencies between and1 Hz (**Figure 3**). Taken together, these two trends are indicative of a viscoelastic material that initially experiences a solid-like state wherein it stiffens to maintain its structure, followed by a more liquid-like state during stress relaxation.) [46]. Additionally, at all measured frequencies, the ratio of G’/G” > 1, which suggests hydrogel-like properties of the LN extracellular tissue (**Figure 3**).

**Figure 3:**
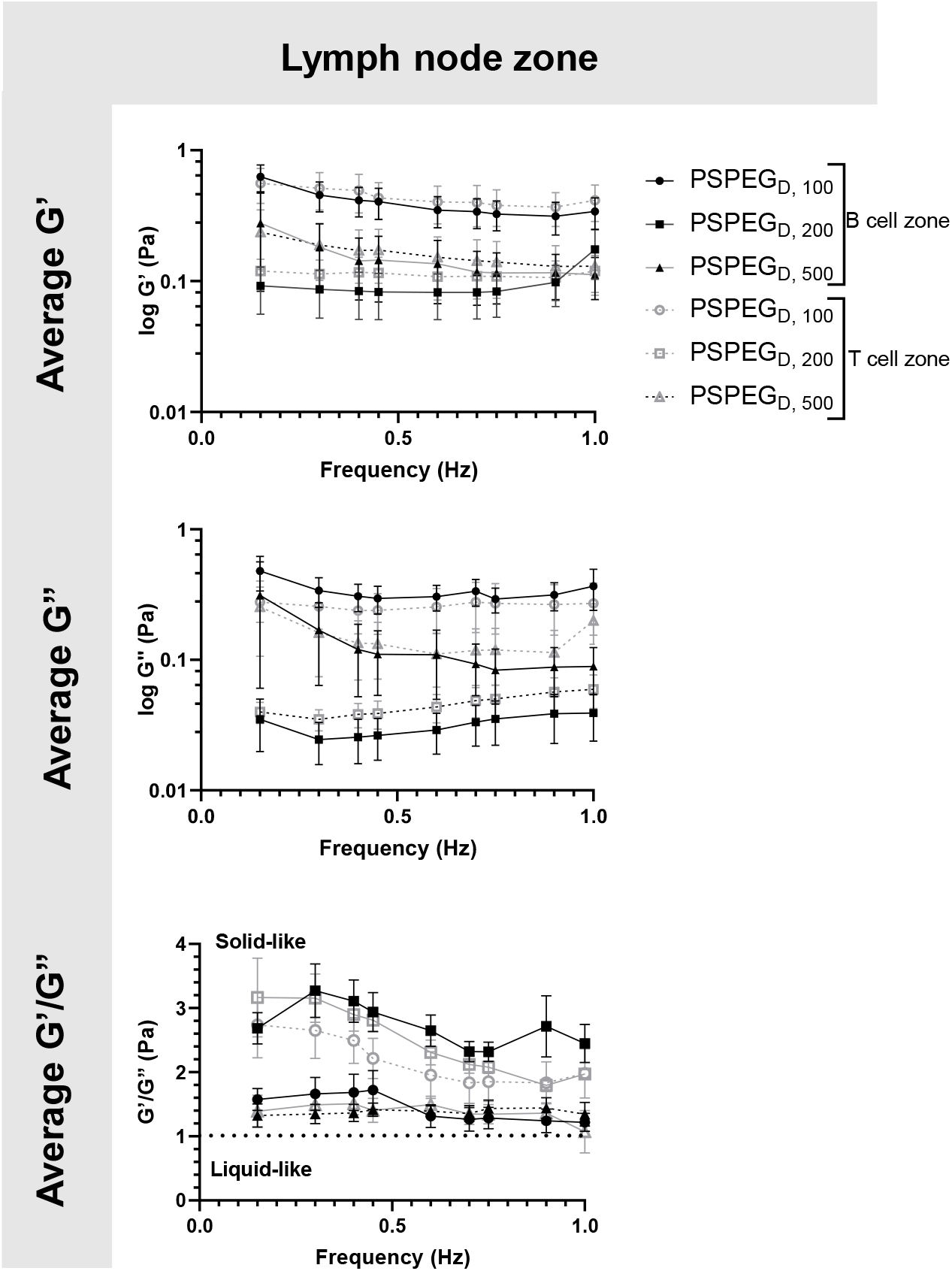
B and T cell zones in LNs exhibit viscoelastic properties. First row shows elastic and second row shows loss modulus of densely PEGylated 100, 200, and 500 nm particles in B/T cell zones and sdLN/mLNs. Last row shows **r**atio of elastic modulus to plastic modulus. Data shown as mean ± SEM (*n* = 5 - 10).

There are several parameters that influence ECM viscoelasticity including composition (collagen, elastin, and other proteins), fiber organization, and types of bonds holding structures together. Dense collagen concentration causes decreased viscoelasticity in tissue and can alter key cellular functions, such as morphogenesis [47]. Cellular interactions also largely affect matrix geometry, which in turn affects matrix stiffness [48, 49]. Cells can cause fibers to align in the direction of load, resulting in a stiffer material [48]. This realignment of fibers allows cells to communicate even at great distances through propagation of mechanical forces [50]. Type and density of crosslinker greatly affects stiffness of tissue and has become a key aspect that researchers tune when creating in vitro ECM models [51, 52]. Weak bonds, such as hydrogen or van der Waals, are overshadowed by strong covalent bonds that can dissipate energy and induce an elastic response [53]. Therefore, the higher the concentration of covalent bonds found within tissue, the stiffer the material and the lower the viscoelasticity. It is likely that the initial response of a solid to a more elastic material is a crucial feature of LNs to accommodate for their potential of rapid expansion and contraction during inflammatory processes [2, 9, 54]. Viscoelastic properties of extracellular tissues may also affect cellular behavior, as it has been shown that mechanical properties of the external environment, including viscosity and viscoelasticity, may affect cell migration and spread [10–12]. In the LNs, cells are particularly motile [55] and there is a constant influx of migrating immune cells entering the LNs from peripheral tissues via lymphatic vessels, as well as immune cells circulating through the LNs of the body via blood vessels [55], making studies of tissue viscoelasticity particularly important.

### Nanoparticles reveal differences in extracellular spaces based on LN location within the body

LNs are found throughout the body, processing antigens and other materials transported via lymphatics from various tissues. We investigated whether LN location, specifically skin draining vs. mesentery (gastrointestinal, GI, draining) LNs (sdLN, mLN), affects the interstitial tissue spaces of the LN. We found that for both sdLN and mLN, dense PEG coatings caused nanoparticles to diffuse through the extracellular tissue space most effectively, as indicated by the highest diffusion coefficient of 4.1 ± 1.3 μm^2^/s, 3.3 ± 0.6 μm^2^/s, and 1.5 ± 0.3 μm^2^/s for 100, 200, and 500 nm PSPEG_D_, respectively, in sdLNs and 2.5 ± 0.6 μm^2^/s, 1.7 ± 0.6 μm^2^/s, and 1.3 ± 0.3 μm^2^/s for 100, 200, and 500 nm PSPEG_D_ in mLNs (**Figure 4A, Supplementary Figure 3**) and proceeded with our analysis of LN extracellular tissue space using these nanoparticles as probes. We found that MSD is consistently slightly lower for mLN compared to sdLN for both 100 nm, 200 nm, and 500 nm PSPEG_D_ (**Figure 4B**), as also illustrated through their representative trajectories (**Figure 4C**). Despite the general trend toward a reduced MSD in mLN, these results are not significant, likely due to the LN-to-LN variation that is illustrated by the variability in diffusion coefficients (**Figure 4A**). We also found that there is a slightly lower pore size for 100, 200, and 500 nm PSPEG_D_ in mLN, with sdLN having a pore size of 890 ± 150 nm, 910 ± 170 nm, and 850 ± 110 nm and mLN having a pore size of 680 ± 100 nm, 770 ± 150 nm, and 890 ± 170 nm, respectively (**Figure 5A**). We determined that the relative frequency of nanoparticles with MSD at 0.5s greater than 0.5 μm^2^ is increased for 100 nm, 200 nm, and 500 nm nanoparticles in the mLN compared to sdLN (**Figure 4D**). Finally, both mLN and sdLN exhibit viscoelastic properties, as indicated by an initial decrease in G’ and G’’ at lower frequencies, followed by an increase in G’ and G’’ at higher frequencies. Additionally, we found that G’/G’’ ratios remained above 1 for microrheological measurements of the sdLN and mLN similar to all other analyses (**Figure 5B**). Altogether our data suggest that LN location may also affect its interstitial tissue properties.

**Figure 4:**
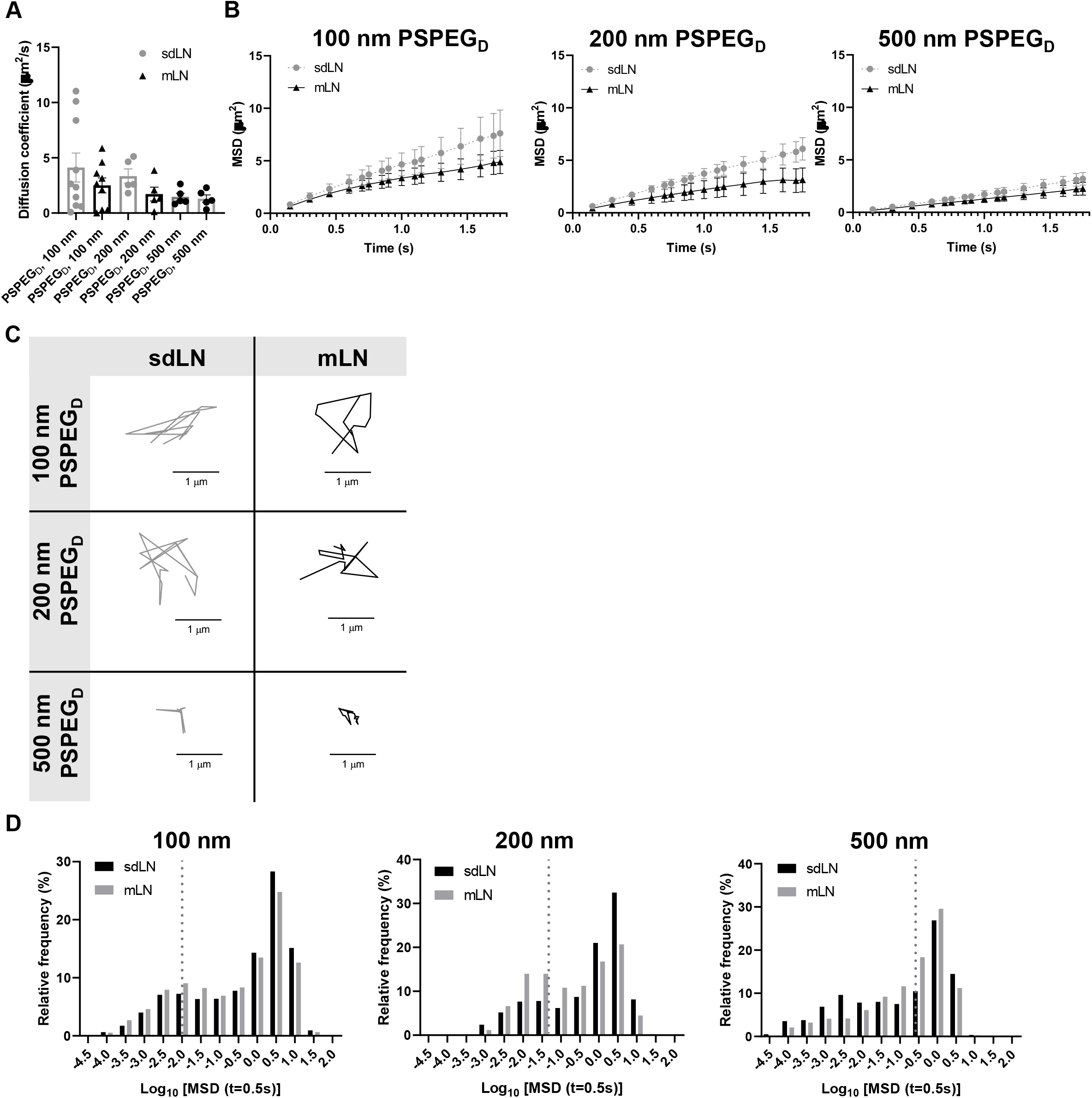
Diffusion of densely PEGylated particles is node-dependent in sdLN and mLNs. **(A)** Diffusion coefficient of densely PEGylated particles in sdLN and mLNs. **(B)** MSD of densely PEGylated particles in sdLN and mLNs. **(C)** Representative trajectories of densely PEGylated particles in sdLN and mLNs. Distance units are in µm. **(D)** Frequency distribution of densely PEGylated particles in sdLN and mLNs. Diffusivity values to the left of the dotted line indicate particles with MSD values less than the particle diameter. Data shown as mean ± SEM (*n* = 5 - 10).

**Figure 5:**
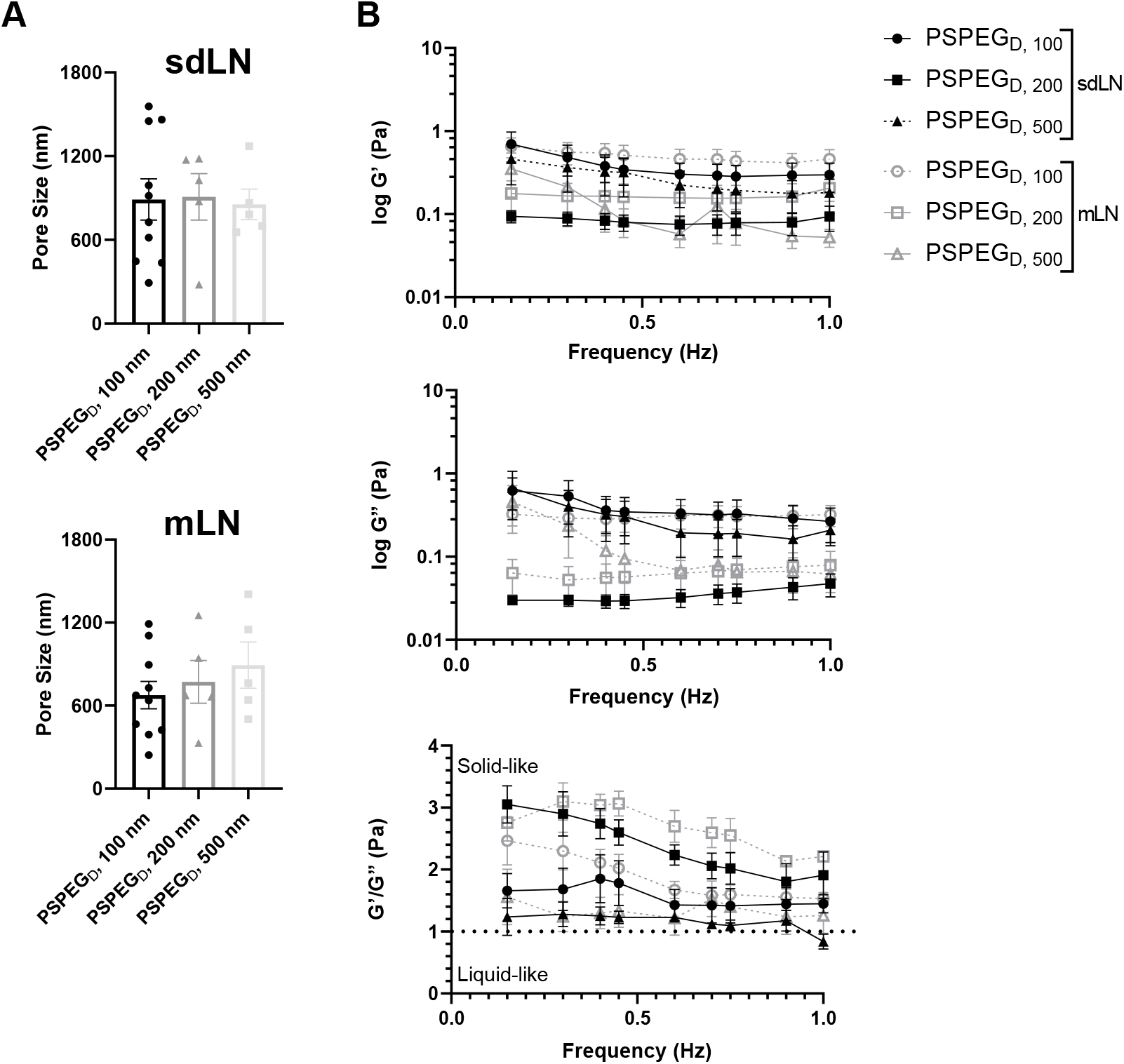
sdLN and mLN show similar pore size and rheological properties. **(A)** Pore size of densely PEGylated particles in sdLN and mLNs. **(B)** Elastic modulus, loss modulus, and ratio of elastic to loss modulus of densely PEGylated 100, 200, and 500 nm particles in sdLN and mLNs. Data shown as mean ± SEM (*n* = 5 - 10).

We hypothesize that differences in extracellular tissue properties of mLN compared to sdLN are likely due to differences in exposure to external materials. LNs at different locations throughout the body are exposed to different stimuli, depending on which peripheral tissue drains into the LN [56, 57]. The keratinized layer on the skin is a significant barrier that effectively keeps pathogenic or foreign materials from infiltrating the underlying dermal tissue, thus preventing the sdLN from being exposed to these materials. In contrast, in the GI tract, lymph draining from the intestinal tissue contains food molecules, such as lipids packaged into chylomicrons, and microbial biproducts from commensal microbes [58]. mLN are thus continuously exposed to potentially inflammatory stimuli, which may explain the differences in their extracellular tissue spacing and properties.

## 4. Conclusions

We have found that nanoparticles densely coated with PEG diffuse through LN interstitial tissue and thus can be used to probe interstitial tissue properties of the LNs. Using this probe, we have demonstrated that LNs are riddled with heterogeneous pores ranging from 100 nm – 1µm. We have also found that nanoparticle diffusion and LN material properties such as viscoelasticity, viscosity, and pore size are not significantly changed in cortex vs. paracortex regions, and that nanoparticle diffusion is reduced in mLN compared to sdLNs. Understanding key measurements including the diffusion coefficient, pore size, viscosity, and viscoelasticity of LN interstitium will allow researchers to better understand how changes in interstitial tissue affect cellular and tissue function, particularly if combined with emerging multi-photon imaging methods. Future work to determine how factors including acute and chronic inflammation or sex affect the interstitial tissue material property-cellular/tissue function axis could give additional insights into how these changes affect cellular functions in the LNs. Finally, our study identifies design criteria for therapeutic nanoparticle delivery within the LN interstitium.

## Author Contributions

Experimental work was done by AR and BM. Data analysis was done by AR, BM, HL, and EV. Writing was done by AR and KM. Review and editing was done by AR and KM.

## Acknowledgements

Financial support from Clark Doctoral Fellowship, University of Maryland (AR) and NIGMS MIRA 1R35GM142835-01(KM).

