## supplementary information for "Multiple particle tracking (MPT) using PEGylated nanoparticles reveals heterogeneity within murine lymph nodes and between lymph nodes at different locations"

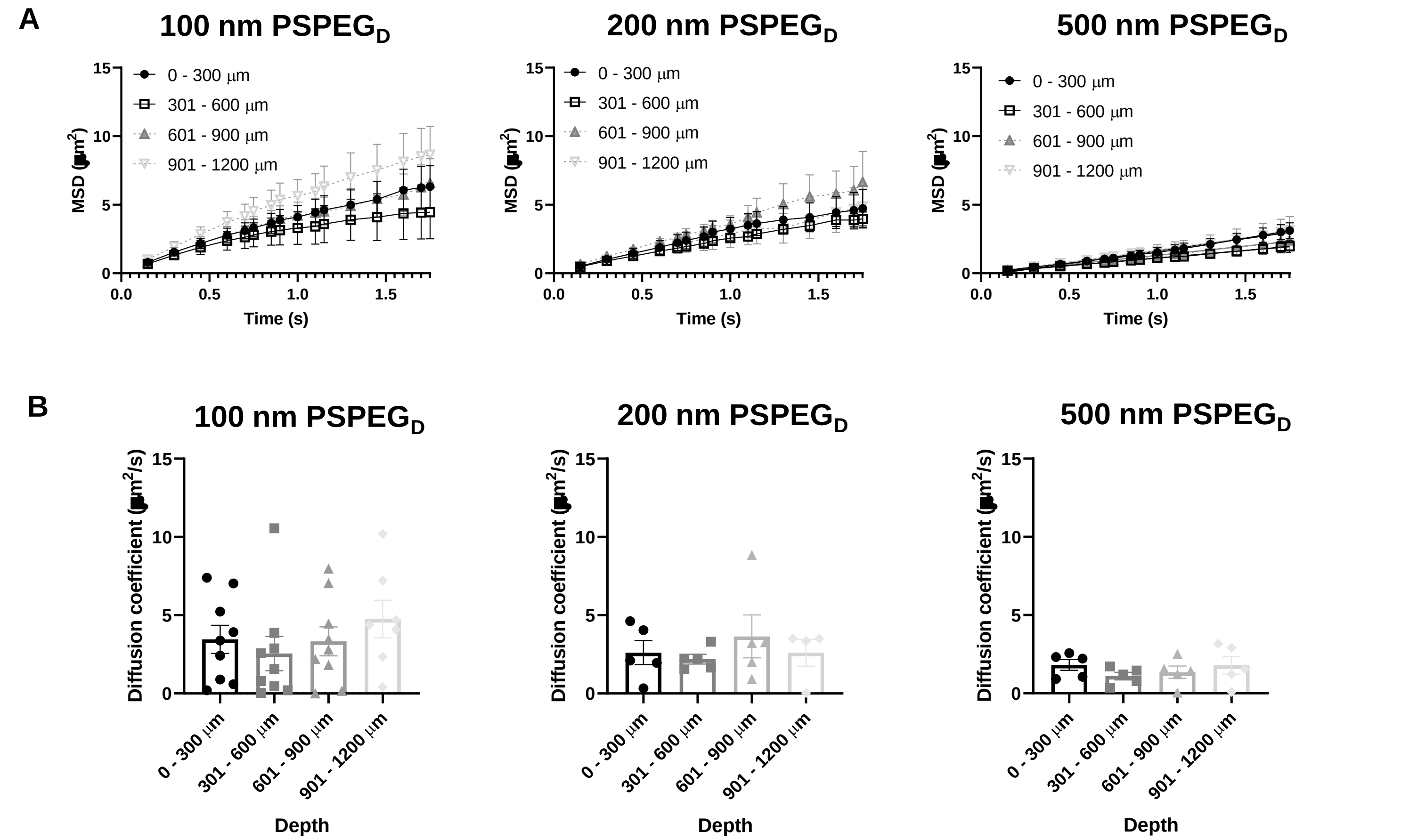


**Supplementary Figure 1:** *Diffusion of densely PEGylated changes throughout sdLN and mLNs.* **(A)** MSD and **(B)** diffusion coefficient of densely PEGylated 100, 200, and 500 nm particles through different depths of the LN. Data shown as mean ± SEM (*n* = 5 - 10).

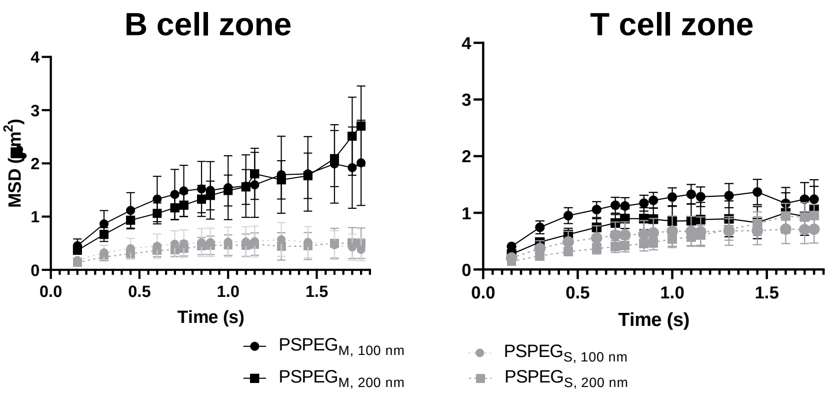


**Supplementary Figure 2:** *Diffusion of PSPEG$_M$ and PS 100 and 200 nm particles in B and T cell zones.* **(A)** MSD of PSPEG$_M$ and PS 100 and 200 nm particles in B cell zones and **(B)** T cell zones reveal that PS particles diffuse less in both areas compared to PSPEG$_M$. Data shown as mean ± SEM (*n* = 5 - 10).

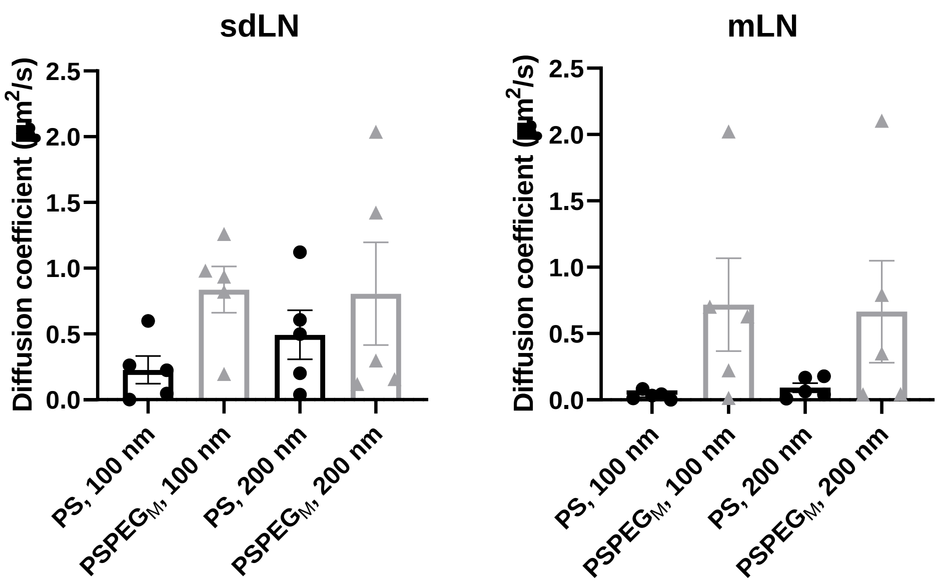


**Supplementary Figure 3:** *Diffusion coefficient of $PSPEG_M$ and PS 100 and 200 nm particles in sdLN and mLNs.* **(A)** Diffusion coefficient of $PSPEG_M$ and PS 100 and 200 nm particles in sdLNs and **(B)** mLNs reveal that PS particles have lower diffusion coefficients regardless of node location. Data shown as mean ± SEM ($n$ = 5 - 10).
